# AlphaBridge: tools for the analysis of predicted biomolecular complexes

**DOI:** 10.1101/2024.10.23.619601

**Authors:** Daniel Álvarez-Salmoral, Razvan Borza, Carmen Maiella, Bjørn P.Y. Kwee, Ren Xie, Robbie P. Joosten, Maarten L. Hekkelman, Anastassis Perrakis

## Abstract

Artificial intelligence (AI)-powered protein structure prediction has transformed how scientists explore macromolecular function. AI-based prediction of macromolecular complexes is increasingly used for evaluating the likelihood of proteins forming complexes with other proteins, nucleic acids, lipids, sugars, or small-molecule ligands. Efficient tools are needed to evaluate these predicted models. We introduce an approach based on combining the confidence metrics of AlphaFold3 to enable clustering of sequence motifs participating in binary interactions and subsequently in 3D interfaces of complexes. Interaction interfaces within confidence limits are finally visualised in 2D using chord diagrams and network graphs. The analysis and visualisation are implemented in a web tool, which links them with interactive graphics and summary tables of predicted interfaces and intermolecular interactions, including confidence scores. Finally, we demonstrate real-life examples of how AlphaBridge is used for providing an efficient way to assess and validate predicted protein complexes and interfaces. The reproducible, objective and automated procedures we present provide a straightforward critical assessment of structure prediction of biomolecular complexes, that should be consulted before conducting more resource-intensive analyses.

## Introduction

Protein structure prediction algorithms have advanced rapidly, extending also into predicting macromolecular complexes. AlphaFold spearheaded this revolution: AlphaFold2^1^ initially showcased unprecedented accuracy in predicting individual protein’s structures. The AlphaFold Protein Structure Database established by Google DeepMind and EMBL-EBI^2^ made these developments accessible to all. Concurrently, ColabFold^3^ offered a fast and accessible solution for predicting protein structures to all scientists through an intuitive user-friendly interface and by leveraging powerful computing resources using efficient search algorithms, matching AlphaFold’s accuracy on critical benchmarks like CASP14^4^ and ClusPro4^5^. Independent approaches notably included RoseTTAFold^6^ and the ESM Metagenomic Atlas^7^.

Extending prediction from individual proteins to the structures of protein complexes, rapidly followed. AlphaFold-Multimer was trained specifically for inputs of known stoichiometry, allowing modelling of protein-protein interactions^8^. Derivative approaches included the AlphaPulldown^9^ idea to use prediction for enabling “virtual pull-down” experiments for validating plausible protein-protein interactions. Furthermore, Predictomes^10^ demonstrated the power of this “virtual pull-down” approach for discovery of new biology, in a closed set of proteins involved in genome maintenance. Importantly, the issue of homo-oligomerization of key proteomes has also been addressed^11^. In addition, AlphaFill ^12^ provided a resource of potential complexes for predicted protein models with about 2,700 ligands that are present in experimental structures. Most recently, AlphaFold3^13^ not only provided algorithmic innovations to the protein structure prediction problem but also allowed efficient and reliable prediction of protein complexes with not only other proteins, but also with nucleic acids and other common biological ligands. These latest AlphaFold developments^13^, notwithstanding licensing issues that preclude “pull-down” screening approaches, were incorporated in a user-friendly web server that popularised the use of protein structure predictions even further, making them available to an ever-growing user base. Similar developments, for example RoseTTAFoldNA^14^ for protein-nucleic acid interactions, or NeuralPLexer^15^ for directly predicting protein–ligand complex structures using ligand molecular graph inputs, are further pushing the boundaries of deep learning applications for studying macromolecular complexes.

Predicted models are nowadays closely coupled to metrics allowing to validate the quality of the prediction, locally and globally. Existing reliability metrics are an excellent aid for experts and can be understood well by keen new users of the technology. The advent of AlphaFold3, with its straightforward availability, speed, and ability to predict protein complexes with proteins, DNA, RNA, as well as select ions, sugars, lipids and biological ligands, draws researchers from different fields into structural biology. Hence, new tools that validate predicted models with intuitive scoring schemes and visualise specifically the reliability and nature of interactions in predicted protein complexes are of essence.

Current approaches to this challenge can be broadly categorized into those that leverage pre-computed confidence metrics, such as the PAE Viewer^16^, and those that perform ab initio assessment based on the predicted three-dimensional structure, such as Pickluster^17^ or DeepUMQA3^18^; concurrent with this manuscript, efforts have been made to systematically catalogue these methodologies^19,20^. Despite this activity, prevailing tools exhibit notable limitations, e.g. many are specialized for protein-protein interactions and lack generalizability to other biomolecular partners. Furthermore, they often fail to provide an automated, integrated, and intuitive output, ultimately relying on the user’s expertise for final interpretation.

AlphaBridge is a comprehensive computational toolkit for the post-processing, detection, analysis, and scoring of interaction interfaces in predicted macromolecular complexes. AlphaBridge provides a unified framework to detect and evaluate interactions across diverse component types. By integrating these analyses into an interactive and intuitive visualization platform, the system makes complex structural data easily interpretable and actionable for researchers across a wide range of scientific fields.

## Results

### Detection of confidently predicted interactions between biomolecules

AlphaFold3 assigns distinct fundamental sequential units (“tokens”) depending on the type of molecule being predicted: a residue for proteins and a nucleotide for DNA or RNA; or each atom for post translational modifications of amino acids or nucleotides (PTMs), ligands, and ions. For the AlphaBridge calculations we keep these definitions; an exception is non-glycan PTMs for which all atoms of the modification are treated as part of the corresponding residue.

For a given biomolecular complex structure model (e.g. replication factor A, a complex between three protein chains, Figure 1A) AlphaFold3 provides three confidence metrics (Figure 1B): the predicted local distance difference test (pLDDT), which measures confidence in the placement of each atom in its local environment; the pairwise aligned error (PAE), quantifying the error in the relative position of two tokens; and the contact probability.

**Figure 1.**
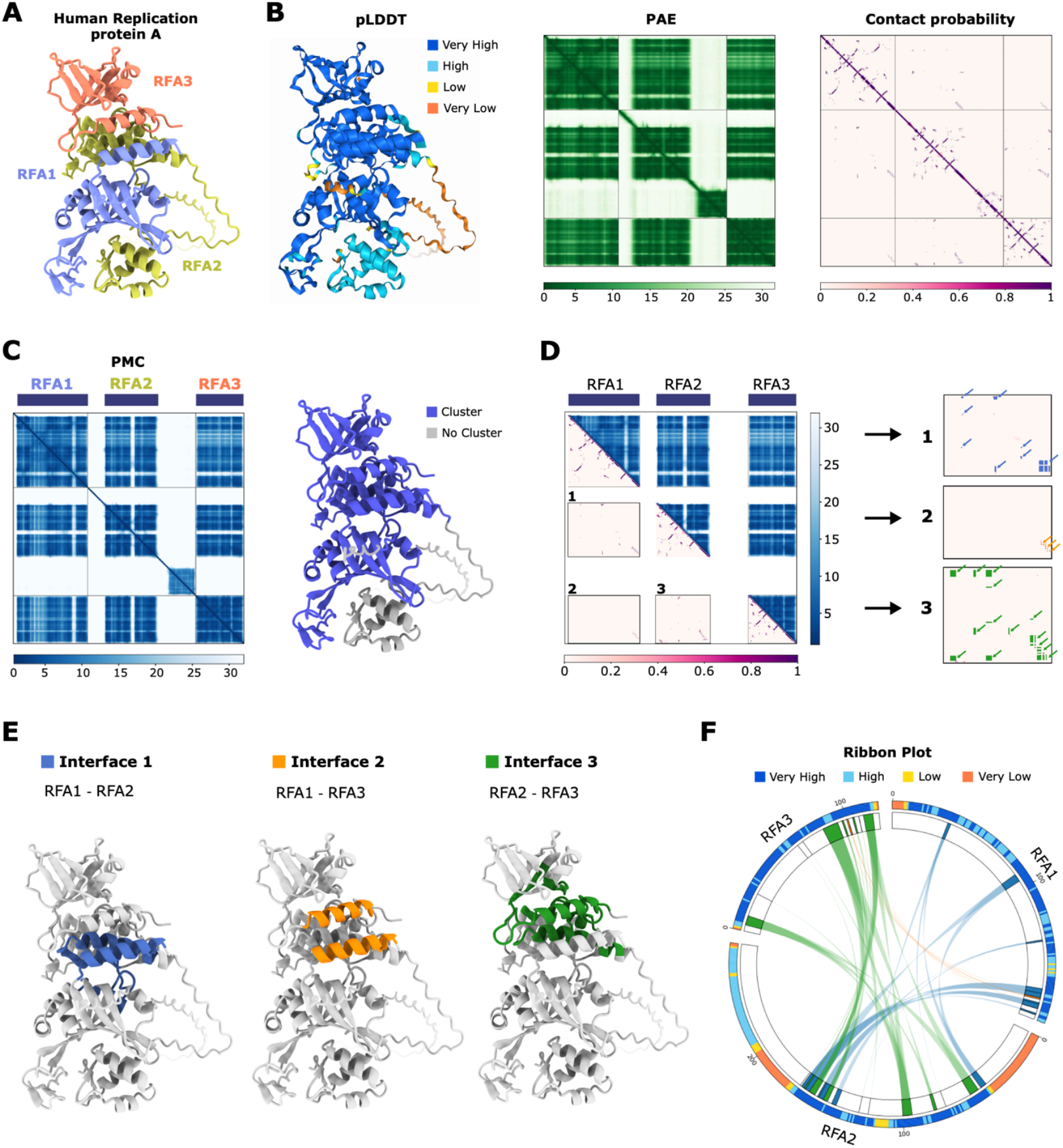
Illustrative case demonstrating Alphabridge analysis of the human replication protein A complex (RPA). **(A)** The AlphaFold3 model of RPA coloured by chain. **(B)** The same model coloured by the pLDDT score (left), the corresponding PAE (middle) and contact probability (right) matrices. **(C)** The Predicted Merged Confidence (PMC) matrix with the regions assigned to the single detected multi-component module annotated by three blue bars on top (left) and the predicted structure with the detected module annotated (right). **(D)** A combination of the PMC matrix (upper right half) and the contact probability matrix (lower left half) with the three interfaces defined between interacting sequential regions between binary combinations of the biomolecules within the multi-component module annotated (left) and the corresponding contact links for each interface (right). **(E)** The three interfaces mapped in the structure of the complex. **(F)** The AlphaBridge diagram for this predicted complex.

We combine the PAE and pLDDT information, in a new matrix we call the Predicted Merged Confidence (PMC) which is used to find “multi-component modules”, non-sequential regions of complex components that are confidently predicted to form a distinct structural unit (Figure 1C). To detect such modules, we implemented a graph-based community clustering approach^21^ inspired by prior work using a similar method for protein domain detection^22^, where the Leiden community clustering algorithm was used. Here, rather than extracting protein domains from a single structure, we combine regions from multiple biomolecules in the predicted biomolecular complex. These multi-component modules are the basic units inside which we proceed to detect the interacting binary interfaces.

To define the binary interfaces, we first mask-out from the contact probability matrix the regions outside the multi-component modules (Figure 1D). Next, we detect ‘contact links’, defined as sequential regions between two biomolecules that have a score higher than a set threshold in the unmasked regions of the contact probability matrix (Figure 1D). For detecting contact links we follow a multidimensional image processing approach, using two-pass connected-component analysis. Sequential contact links that are less than three residues (or nucleotides) apart, are combined to a single link. The collection of all contact links between two components of each multi-component module, forms an interacting binary interface.

This process is iterated for all multi-component modules, until all binary interfaces are characterised (Figure 1E). This way, for each biomolecular complex, we define several interacting interfaces, and for each interacting interface we define contact links. The number of contact links and interfaces can vary depending on the cut-off value used for masking the contact probability matrix. Varying this value can increase or decrease the confidence with which contacts and interfaces are assigned.

We first discuss how this evaluation procedure is used to visualise confidently predicted interfaces to help expert and non-expert users to better evaluate AlphaFold predictions. Then, we will show how we use it to calculate confidence scores for the interaction of individual interfaces, between biomolecules, as well as global scores for a predicted complex, offering a complementary criterion for scoring and ranking the quality of predicted interactions.

### Construction of the AlphaBridge diagrams for biomolecular complexes

The assigned interfaces and contact links are then shown in a circular layout^23^ as a chord diagram (Figure 1F). Each chain of amino-acids or nucleotides occupies the outer arc in a manner proportional to its sequence length. Non-glycan PTMs are annotated in their corresponding position. Each ligand or glycan PTM occupies an arc of 10^°^in the outer ring of the diagram, regardless of its size in atoms; ions occupy 5^°^. If the sum for all PTMs, ligands, and ions exceeds 90^°^, the total arc to present them is limited to 90^°^and each individual arc is adjusted. This adjustment prioritizes the display of the polymer components (proteins and nucleic acids) involved in the interactions.

Interfaces are depicted as sets of curves, or “bridges”, between the regions that make up contact links, with different colours for each interface. The outer ring displays the pLDDT for each chain as a thick arc. This type of visualisation integrates various complementary information sources into a single, easily interpretable visualisation allowing users to quickly assess the number and confidence of all interactions and the number of interfaces in each complex.

### Network-based representation for biomolecular complexes

We calculate two types of network representations, based on the identified interfaces. The simplest of the two is the “biomolecular network”, an undirected graph in which each node represents a biomolecule, and each edge denotes that at least one interface is detected between these two nodes. This representation provides a high-level view of the interactions, like networks from experimental data (e.g. STRING) and emphasizes global connectivity between biomolecules. A more complex presentation is the “interface network”, a multigraph that offers a finer-grained perspective by representing sequential regions of contact links. Nodes represent “merged” contact links from interfaces: contact links are merged if they have at least one residue on common. Edges are used to connect the regions within the biomolecule, and to present the connection of each of these regions to regions of other biomolecules of the complex. This dual-edge structure allows for a more detailed examination of interaction interfaces, while preserving contextual sequence information.

### Scoring the confidence of biomolecular complexes and individual interaction interfaces

Assignment of the interaction interfaces between subunits, offers an opportunity to develop scores that specifically consider this new information, both at the entire structure, but also focusing on individual interaction interfaces. We provide two new scores to help users assess the likelihood of a predicted complex.

The “*predicted interaction Confidence Score*” (*piCS*) is based on the PMC matrix, weighing favourably complexes with confident predictions for both the local and the relative structure of all biomolecules. This score carries inherently the possibility of being able to be compute for each individual interface, allowing to assign scores that indicate how important each interface is for complex formation. It should be noted that the global scores exclude tokens corresponding to ligands, ions, or glycans in the calculation, while the individual interface scores consider interactions with all forms of ligands. Details on the calculations are available in Methods.

The *AlphaBridge* score is based on the cut-off value used for masking the contact probability matrix before interfaces are detected, a key parameter during the process of evaluating the interaction interfaces we described above. This cut-off value is a user-selected parameter, with three “pre-set” different cutoff settings: “high confidence” (0.9), “default” (0.7) and “low confidence” (0.5). It can be used as a “visual scoring” criterion, helping to intuitively understand the confidence of predicted interactions by toggling between settings. It is also easily accessible programmatically for higher-throughput applications. Varying this cut-off value allows to compute the level above which an interface between two components of the complex is detected, and that information is then used to assign scores in two levels: i) for each pair of biomolecules, as the maximum value of piCS for all interfaces contributing to this association; and ii) as a global score, as a weighted average between all binary biomolecular interactions. Details on the calculations are available in Methods.

Note, that the *piCS* and the *AlphaBridge* score are in a range of 0 (not confident) to 1 (confident) similar to the *iPTM* score that is used by AlphaFold.

### The AlphaBridge webserver

Online analysis of AlphaFold3 results can be performed by uploading the AlphaFold3 output “*zip*” file to https://alpha-bridge.eu. All information is processed automatically and an interactive AlphaBridge diagram is constructed to the left, while an interactive 3D structure viewer is shown to the right (Figure 2A). Users can choose between three confidence levels for interface detection: low, default, and high as discussed above. Toggling between these settings is fast, allowing “visual scoring” for the confidence of individual interactions. The *ipTM* score for the complex prediction confidence from AlphaFold3 is displayed alongside the global *piCS* and the *AlphaBridge* scores. The AlphaBridge diagram and the structure viewport are connected, in such a way that clicking on a contact link, displays that interface. The outer ring of the AlphaBridge diagram shows the per-chain colours that are also used for colouring in the interactive graphics; mouse over the outer ring displays pLDDT values and residue information; post-translationally modified residues are designated by a line between the two outer rings of the AlphaBridge diagram. Contact links belonging to the same interface have the same colour; mouse-over on contact links displays the interface number and the residues involved. The AlphaBridge diagram can be downloaded as an SVG file for use in presentations or publications. In the viewport, the user can toggle between structure colouring by chain, by pLDDT, or by interface (Figure 2B,C); snapshots can be taken with the integrated functionality of the viewer. A collapsible table shows the interface contact link details for each interface (Figure 2D), including the *piCS* value for each individual interface next to its name. The table is interactively linked to the 3D viewport, thus clicking on an interface or a specific contact link focuses on the relevant residues in the 3D viewer. The *biomolecular network* is shown by default; the colours of each node are the same as for the AlphaBridge diagram and edges are black (Figure 2E). The numbers in each edge correspond to the *AlphaBridge* score for each binary interaction and can be used to judge the relative confidence between different binary interactions. Users can toggle between the default *biomolecular network* and the more complex *interface network*, which shows the inter-connections of sequential regions between biomolecules (same colour as the interface), as well as the intra-connection between nodes that belong to the same biomolecule (same colour as the nodes for this biomolecule).Finally, the PAE, contact probability and PMC matrices are displayed at the end of the web page, for reference.

**Figure 2.**
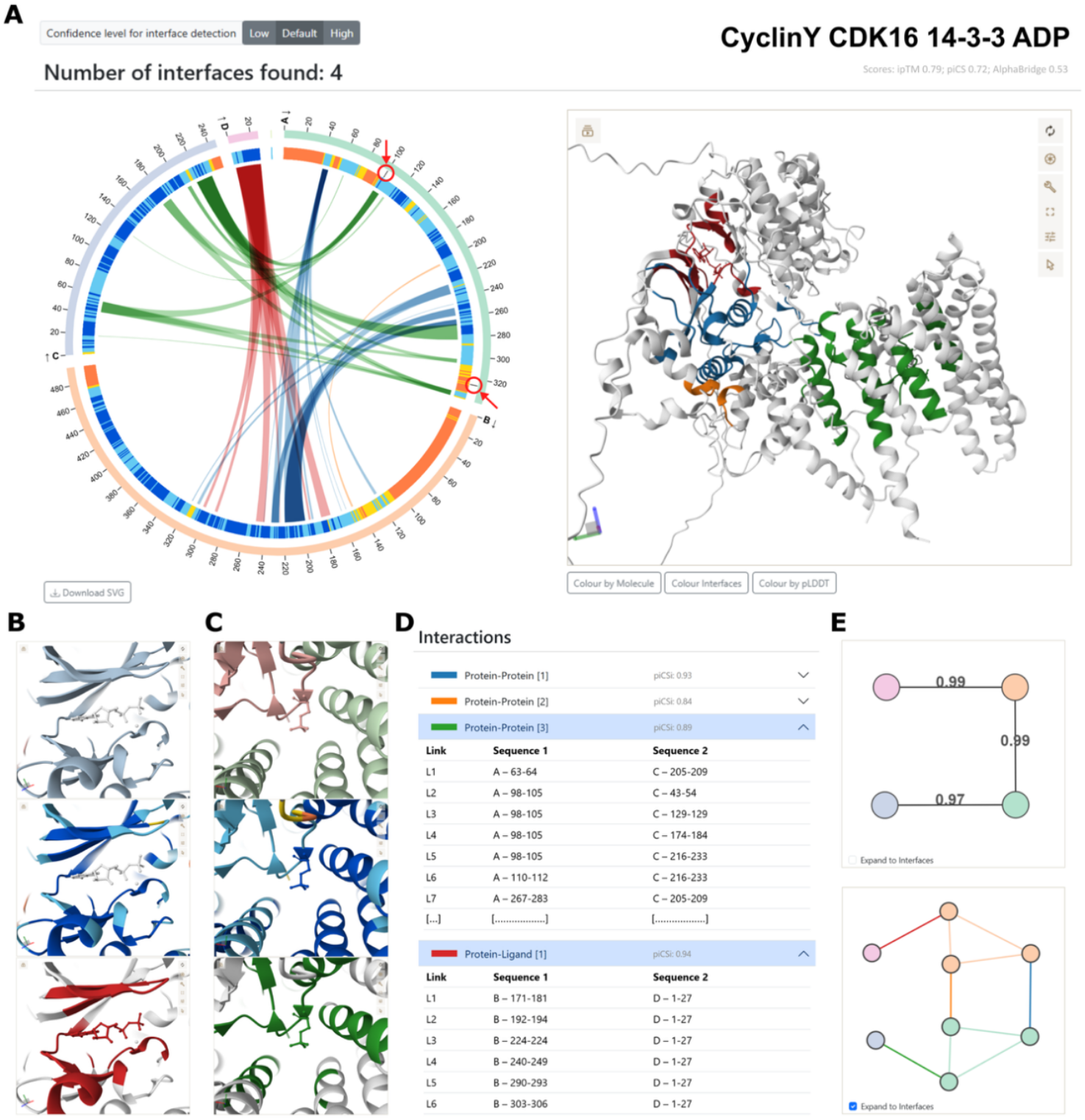
Screenshots from AlphaBridge webserver analysis for a structure prediction of the Cyclin Y - CDK16 - 14-3-3 - ADP complex. **(A)** The AlphaBridge diagram to the left and the interactive 3D-view to the right; arrows have been added to highlight two modelled PTMs that are important for interaction24. **(B)** The 3D-viewport zoomed into that ADP-binding interface, coloured by molecule, pLDDT value and by interface. **(C)** The same for the interface between CyclinY:14-3-3, highlighting the phosphoserine PTM. **(D)** The table of all interactions by interface, with details on the CyclinY:14-3-3 interface highlighting the involvement of a PTM, and the CDK16:ADP interface shown in more detail. **(E)** The *biomolecular network* (top) with the *AlphaBridge* scores along the edges, and the *interface network* (bottom).

### Practical examples of using AlphaBridge

Condensin I and II facilitate the spatial rearrangement of the human genome upon mitotic entry. Recently, others and we, identified MIS18-binding protein 1 (M18BP1) as a long-sought bridge for condensin II localization to chromatin ^25^. AlphaFold3 models and AlphaBridge diagrams (Figure 3A) clearly show that M18BP1 directly binds the CAP-G2 subunit, like MCPH1, a known condensin II antagonist. The AlphaBridge diagrams, clearly show that binding is on the same site of the CAP-G2 subunit. The AlphaBridge scores for the MCPH1:CAP-G2 (0.83) and M18BP1:CAP-G2 (0.85) interactions, suggest a competitive regulatory mechanism for condensin II localization^25^.

**Figure 3.**
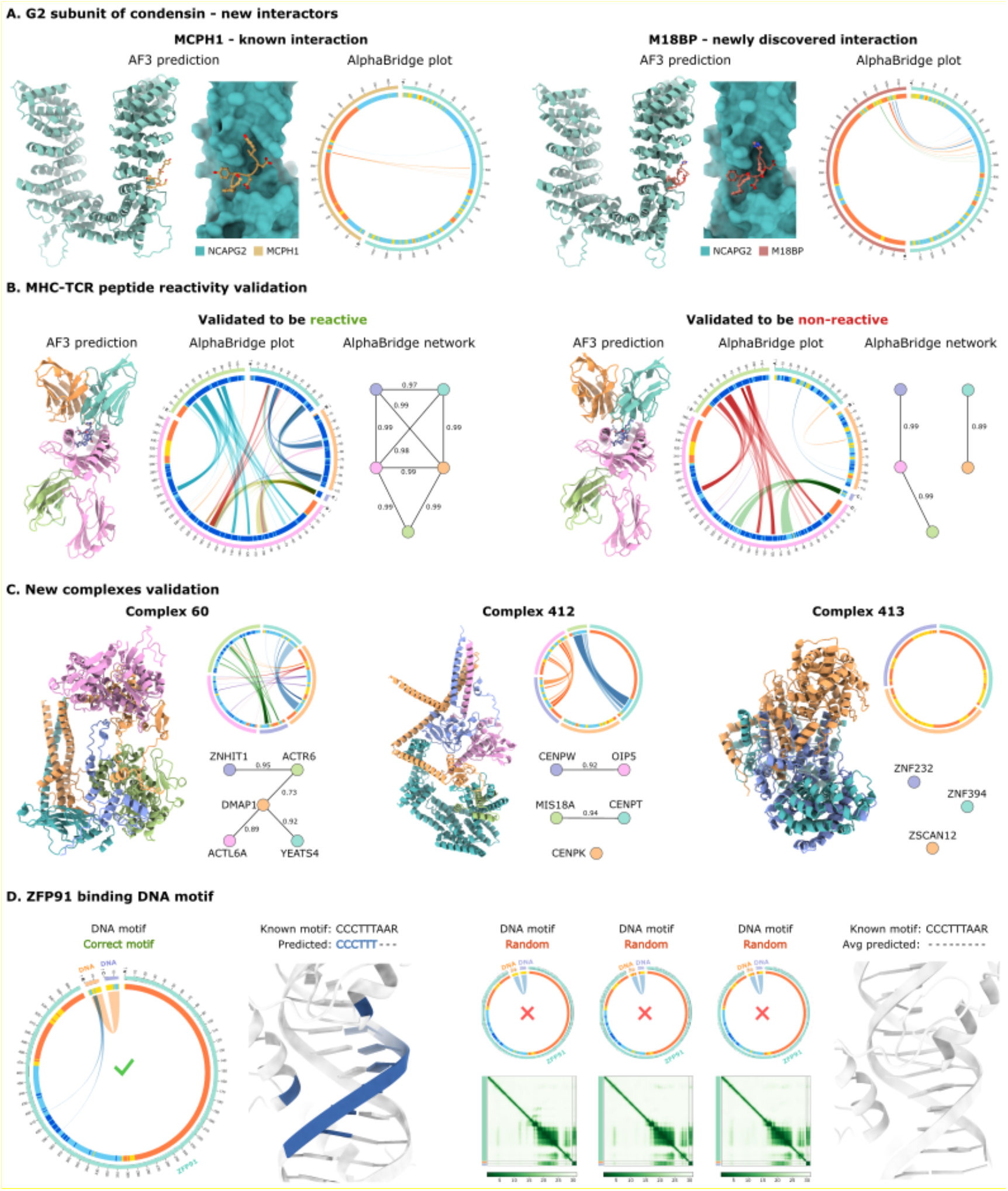
Representative predicted biological assemblies, demonstrating AlphaBridge’s assessment of confident predicted interactions. **(A)** Comparison of a known interaction (CAPG2–MCPH1; left) and a newly discovered interaction (CAPG2–M18BP; right); for each prediction we show (from left to right) the mode predicted by AlphaFold3 coloured by chain, the AlphaBridge diagram, and a zoom-in the interaction site, to show the antagonistic binding mode **(B)** Comparison of validated to be reactive (left) and validated to be non-reactive (right) TCR:pMHC complexes: shown from left to right are the predicted model, the AlphaBridge diagram, and the biomolecular network; the non-reactive complex is clearly distinguishable visually in the AlphaBridge diagram and the biomolecular network **(C)** Multi-protein complexes from the AHMPC dataset: for each example, the predicted structure is shown coloured by chain (left), with the corresponding AlphaBridge diagram (upper right) and biomolecular network (lower right) illustrating whether subunits interact stably (Complex 60; left) or should be refined (Complex 412; middle) or discarded (Complex 413; right). **(D)** Validation of ZFP91 binding to its cognate DNA motif: to the left the AlphaBridge diagram for the correct motif and the predicted structure (the identified interface highlighted in blue); to the right the AlphaBridge diagrams (top) and PAE matrices (bottom) for three random motifs showing non-significant binding, and a predicted structure with a random motif.

The T cell-mediated adaptive immune response relies on a set of diverse T cell receptors (TCRs) that recognize peptides (p) presented by major histocompatibility complex (MHC) molecules. Predicting which TCR:pMHC interactions may elicit an immune response remains a fundamental challenge. Significant efforts, focused on improving the quality of experimental datasets of TCR:pMHC interactions, are now enabling differentiating between functionally reactive and non-reactive pairs of TCR-pMHCs for hundreds of TCRs and specific antigenic peptides (epitopes)^26^. While the structures of AlphaFold3 predictions of reactive and non-reactive complexes (Figure 3B) look similar, there is enough information in the validation metrics to allow AlphaBridge diagram and the biomolecular network to visually differentiate between reactive (left) and non-reactive cases (right). The AlphaBridge score has been specifically modified to quantify the TCR-pMHC interactions (TCRbridge) and shows good promise on differentiating reactive and non-reactive TCR – epitope pairs^26^.

In an ongoing project, our lab developed a comprehensive resource of newly predicted human multi-protein complexes referred to as AHMPC^27^. These complexes are inferred from large-scale structural modelling of binary protein–protein interactions across the human proteome^28^ and further supported by orthogonal functional genomics data to enhance their biological relevance^29^. AlphaBridge diagrams and biomolecular networks (Figure 3C) provide an intuitive way to validate, refine, or if necessary, discard components of the predicted complexes. AlphaBridge scores allow a very easy filter for what should be considered a confident interaction within the assembly.

Our lab recently described Zincore^30^, a protein complex that locks specific zinc finger transcription factors (ZNFs), including ZFP91, onto DNA. As part of our effort to understand ZFP91 binding to DNA, prior to structure determination by cryo-EM (PDB:9HJT) we made AlphaFold3 predictions of ZFP91 towards different DNA motifs. While model inspection shows no discernible differences in ZFP91 binding between cognate target DNA motifs and random sequences (Figure 3D), AlphaBridge diagrams visually differentiate between correct and random motifs, suggesting the use of AlphaBridge as a valuable tool recognize possible new binding target motifs for DNA binding proteins, based on AlphaFold3 modelling.

## Discussion

AlphaBridge is designed to aid all scientists who use structure prediction of macromolecular complexes to validate and interpret the predicted models. First, it provides an automated, reproducible, and reliable algorithm to detect linear sequence motifs (contact links) involved in binary interfaces between predicted structures of macromolecules, considering the metrics that describe the reliability of the predictions. And second, it offers an interactive web interface connecting an intuitive way to present interactions in “AlphaBridge diagrams” (useful for presentations and publications) and network graphs, together with live 3D visualisation (suitable for interactive analysis).

The PMC matrix introduced here provides a reliable, accessible, and reproducible method for distinguishing biologically meaningful interactions, extending beyond conventional analysis of pLDDT and PAE matrices. Additionally, it offers a visualization tool tailored for specialized applications. The PMC and contact probability matrixes, allow the delineation of the linear motifs that make the contact links, enabling straightforward inspection of regions of interaction for non-experts. The detection of these contacts using multiple cut-off values as threshold, offered the opportunity to define a user-accessible way to detect less or more confident interactions.

Another opportunity offered by defining interfaces between any component of a macromolecular complex in a chemistry-agnostic way, is in scoring the importance of various interactions. The *ipTM* score is well-established and useful for assessing the predicted overall accuracy of macromolecular complexes, as well as the accuracy of the prediction of complexes between individual subunits. However, it has its shortcoming, and alternative scores that are better suited for complex validation have been proposed^31^.

The scores we present here (piCS and AlphaBridge score) have the advantage that they are inherently computable for individual interfaces, not only between subunits. This enables users to assess the contribution of specific regions to complex formation, particularly in multicomponent assemblies. Namely, piCS builds on the PMC matrix, stronger weighing well-predicted structured interfaces. It must be noted that while such information is clearly present in the PAE matrix, as well as in the contact probability matrixes and inter-chain *ipTM* matrixes that are available in the extended AlphaFold3 output, here we consolidate that information and present it to the end-user in a more intuitive and interactive manner. In addition, we introduce the AlphaBridge score, which takes advantage of the procedure for assigning interfaces with specific confidence intervals, to provide a score for the likelihood of interaction between specific biomolecules in the complex. The TCRbridge score ^26^ we briefly discussed above (Figure 3B), is proving successful for distinguishing between reactive versus non-reactive pairs of TCR and antigens presented by MHC.

The circular layout visualisation we employ, is familiar to many users (as it has been used for instance to show interactions derived from cross-link proteomics applications^32^), has the possibility to represent increasingly complex data in two dimensions^33^, and can also be used to integrate additional information. For example, the display of AlphaMissense^34^ predictions (for human proteins) and sequence conservation data (for all proteins) could be considered. Including these additional layers of information, can help analyse and understand the functional implications of interactions, identify functionally important residues, or indicate potential disease-associated variants involved in interfaces, enriching the biological relevance of the interactions.

Similarly, the biomolecular graph is familiar to most users and offers a quick overview of interactions. Displaying the scores between each binary biomolecular interaction, also facilitates easy inspection of multi-components complexes. Adopting this scoring in evaluation pipelines, as we show through the examples of the AHMPC dataset ^27^ (Figure 3C), can help to validate if all components of a hypothetical complex interact with each other.

Currently, the AlphaBridge web server works with results from AlphaFold3^13^. It can process predictions that include complexes between proteins and nucleic acids (DNA and RNA), can handle post-translational modifications (PTMs) in proteins, and biological ligands. Our algorithm can be both integrated in additional tools (e.g. OpenFold^35^, ColabFold^3^, Predictomes^10^, or Boltz-2^36^) or straightforwardly enhanced to process output from such servers.

In conclusion, AlphaBridge enables the annotation, interpretation and validation of AlphaFold predictions of macromolecular complexes, aspiring to specifically engage and help non-experts.

## Supporting information

NONE

## Acknowledgements

We are grateful to all the AlphaBridge “beta users” from the NKI and beyond for their valuable feedback, insightful discussions, and enthusiastic support during development. We thank Ida de Vries, Marius Messemaker, Titia Sixma, Thijn Brummelkamp, Wouter Schepper, and Ton Schumacher for their critical reading of the manuscript and their constructive suggestions. Finally, we acknowledge the Research High Performance Computing facility of the Netherlands Cancer Institute for providing and maintaining essential computational resources.

## Methods

### Consideration of state-of-the-art model confidence metrics

For a given biomolecular structure model, AlphaFold3 provides three confidence metrics: i. the per-atom local confidence (pLDDT), which measures confidence in the local structure on a scale from 100 (confident) to 0 (not confident); ii. the pairwise aligned error (PAE), quantifying the error margin in the relative position of two tokens within the predicted structure, on a scale from 0 (confident) to 33 (not confident); and the pairwise token-token contact confidence (contact probability), estimating the confidence that two relative tokens are in contact (i.e. having a distance of 8 Å or less between the representative atoms for each token), on a scale from 1 (confident) to 0 (not confident).

The PAE serves as an effective metric for assessing the confidence in domain packing and the accuracy of the relative domain placement within the predicted structure. However, it is also important to consider the confidence in the local structure by the pLDDT value, estimating how well the prediction would agree with an experimental structure. Variations in the pLDDT score across the macromolecular structure highlight which parts of the predicted model are more reliable and which are less so. Such variations might signify regions of high flexibility or intrinsic disorder, where the structure is either ill-defined or lacks sufficient information for a confident prediction.

Another limitation arises while assessing the PAE value of two related tokens without considering whether these residues engage in meaningful interactions or belong to the same functional site, which poses a notable constraint. It is possible to have high confidence that two residues are far apart within the structure. Lower PAE scores suggest higher confidence in the relative position of two tokens within the predicted structure. However, it does not always infer whether those residues are in close contact or interacting with each other.

These limitations become especially significant when trying to understand the functional implications of residue interactions within a predicted macromolecular complex. To address this, we developed a reinterpretation and visualisation pipeline that compensates for the limitations of relying solely on pLDDT and PAE values.

### Calculating the PMC matrix

In the PMC matrix each token is represented similarly to the PAE. To merge the per atom pLDDT scores with the pairwise PAE token-token matrix confidence scores and obtain the PMC score, we need to first calculate a pseudo-pLDDT (*ψ*_*pLDDT*_) score over all atoms (N) in the pair of tokens *i, j*:

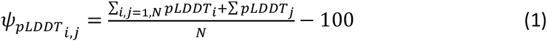

Note, that the vales of *ψ*_*pLDDT*_ are reversed, with the highest score (0) denoting confidence, and the lowest score (−100) denoting lack of confidence, for consistency with the scale of the PAE matrix; then the PMC matrix is calculated as:

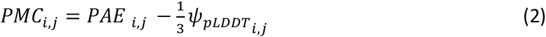

Note that the way that the PMC matrix is calculated, its values are defined on a scale from 0 (confident) to 66 (not confident), similar to the PAE. A special case for calculating the PMC score occurs for pairs involving non-glycan PTMs. These are tokenized per-atom in the PAE matrix, but we consider them as part of their corresponding residue. When calculating the PMC value between a non-glycan PTM residue and any other residue, the minimum PAE value is selected as the representative for that pairwise interaction.

### Defining intra-protein structural modules

The graph-based community clustering approach ^21^ we use is based on the work of T. Croll ^22^. This was originally implemented to detect protein domains in a single sequence and the PAE matrix; here it is modified to work with multiple sequences and the PMC matrix. The algorithm is based upon the Leiden community clustering algorithm ^21^, to identify groups of nodes that are more densely connected to each other than to the rest of the network, greedily optimising modularity in the content of the division of a network into clusters.

By considering also the pLDDT scores, we aim to eliminate the interfaces with a low confidence of the local structure from the pool of potential candidates of interacting interfaces. Rather than extracting protein domains from a single monomer structure, the adapted methodology offers a compilation of distinct structural units from multiple biomolecules (multi-component modules) in the predicted biomolecular complex that share the same biological context within the predicted structure. These multi-component modules are the basic units inside which we proceed to detect the interacting binary interfaces.

### Assigning interfaces

Given an assigned multi-component module, each binary biomolecular combination is considered within the predicted macromolecular complex. By evaluating this subset of interactions, the search space is significantly narrowed down from considering all potential binary interactions across all entities in the predicted structure, to focusing on each specific binary interaction inside the multi-component modules. This approach aims to improve both the accuracy of identifying interacting interfaces and the efficiency of computational processing.

For a given binary biomolecular interaction, all the contact probability scores were extracted within the range of tokens involved in the interaction. A similar approach as described earlier for the PMC matrix is applied for non-glycan PTMs; here the maximum value of the involved tokens is considered as the representative for all pairwise interactions involving the modified residue within the contact probability matrix. A multidimensional image processing approach is applied to detect sequential regions of connected pairwise tokens or ‘contact links’ using a two-pass connected-component analysis. We detect ‘contact links’, that have a score higher than a set threshold in the unmasked regions of the contact probability matrix.

For this approach a graph is created using the contact probability matrix as input. Each vertex holds the necessary information for the comparison heuristically, while the edges represent connections to neighbouring vertices. An algorithm navigates through the graph, assigning labels to the vertices based on their connectivity and the relative values of their neighbours. This connected-component analysis uses the scipy.ndimage module from the SciPy package^37^.

Within a binary interaction, an interacting interface is defined by all shared contact links between two distinct biomolecular entities. Sequential contact links that are less than two residues (or nucleotides) apart, are combined to a single link. The collection of all shared contact links between two components of each multi-component module, forms an interacting binary interface. If a binary interaction contains *n* distinct groups of non-shared contact links, *n* interacting interfaces will be defined for that specific binary interaction.

### Computing the Predicted Interaction Confidence Score (piCS)

To compute the Predicted Interaction Confidence Score (piCS), we first define the set of all values of the contact probability matrix between pairs of tokens that belong to detected multi-component structural modules, *CPMod*.

Then we define the set that excludes all zero values:

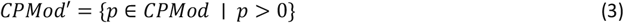

Then we determine the 75th percentile, i.e. the 3rd quartile *Q*_3_(*CPMod′*) and define a new set:

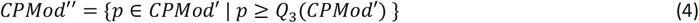

Then we define the contact probability, *CP*, as:

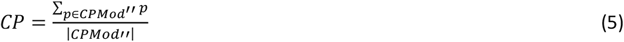

To calculate the piCS score, we first normalise the set of values in the PMC matrix (*PMC)*, to be in a range from 0 (not confident) to 1 (confident), for consistency with the previously defined scores:

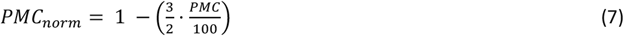

Then we define the average confidence of the normalised set of values, *C*_*PMC*_ as:

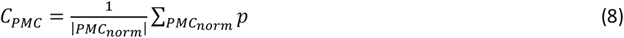

Then the *piCS* score is defined as:

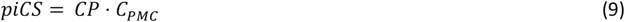

### Computing the AlphaBridge score

The AlphaBridge score quantifies the confidence of detecting molecular interfaces within a complex, based on a thresholding approach applied to the contact probability matrix.

Let τ ∈ [0,1] be the threshold parameter used to mask the contact probability matrix before interface detection. As *τ* ***i***s decreased from its maximum value (τ = 1, where no interfaces are detected) to a user-defined minimum (e.g., τ = 0.5), interfaces begin to emerge. Let k be the total number of interfaces detected above a particular minimum cut-off value for τ. Then, we define 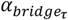 as the threshold value above which each interface *i=[1,k]* is detected.

Next, given a set of biomolecular components of a biomolecular complex, *B* ***=*** *{B*_*1*_, *B*_*2*_, …, *B*_*n*_*}*, we consider all possible binary pairs *(B*_*i*_, *B*_*j*_) within it. For each binary interaction *BI*_*ij*_ where at least one interface has been detected between *B*_*ij*_ and *B*_*j*_, we collect the corresponding 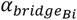 values from their constituent interfaces, and define:

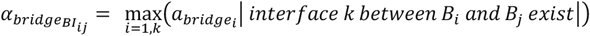

Having defined the 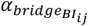 values between all pairs of biomolecules that have at least one interface, last, we define a global score for the entire macromolecular complex, as:

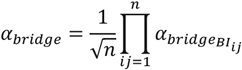

### Network-based representation for biomolecular complexes

The two hierarchically related network representations we developed, the *biomolecular network*, and the *interface network*, both derive from the same underlying confident interactions, but serve different levels of granularity.

The *biomolecular network* is an undirected graph *GP=(VP, EP)* in which nodes (*VP*) correspond to biomolecules within the prediction, and edges (*EP*) indicate confident interactions between pairs of biomolecules. To construct this network, when two biomolecules are connected by one or more interfaces, an edge is drawn to represent the interaction in the network.

The *interface network* is a multigraph *GI=(VI, EI, D)* that offers a finer-grained perspective by representing sequential regions s involved in contact links. Contact links were grouped by interface, allowing each contact link and its corresponding residue range to be associated with a specific interface region. The interface-level interactions are then mapped onto the graph. Nodes (*VI)* in this network correspond to these interaction regions, while edges (*EI)* are divided into two distinct sets: the first set captures intra-connections within a single biomolecule, and the second set represents inter-connections formed by contact links between different biomolecules. This dual-edge structure allows for a detailed examination of interaction interfaces while preserving contextual sequence information.

### Plotting diagrams

The Circos-like layout figures were created using the pyCirclize package^38^, designed for plotting circular figures like Circos Plots and Chord Diagrams in Python. pyCirclize employs a circular layout with sectors and tracks, allowing different data types to be assigned to each sector. Multiple tracks for data plotting can be freely arranged within each sector.

### Implementation and Code availability

The code was developed in Python 3.9 with Anaconda as the package manager to ensure a consistent and reproducible environment. Key modules used include SciPy for performing the two-pass connected-component analysis and PyCirclize for creating circular layout diagrams. The complete code, along with setup instructions and environment configuration, is available in the public repository https://github.com/PDB-REDO/AlphaBridge, which includes detailed guidance on installation and usage.

### Web server implementation

The server was built using the libzeep library^39^, which provides essential tools for HTTP server management, HTML templating, and various components for web server construction in C++. To handle mmCIF files containing the macromolecular model coordinates we integrated libcif++^40^. For model visualization, we used Mol*^41^ as an interactive web component embedded directly into the webpage. To manage JavaScript dependencies, we employed Yarn, an open-source package manager. For interactive data visualisation, specifically for generating circular layout diagrams and network-based representation, we utilised the free and open-source library D3.js.

